# Combining non-linear and spectral cardiac variability metrics differentiates rat alcohol drinking with and without challenge

**DOI:** 10.64898/2026.11.11.765942

**Authors:** Thatiane De Oliveira Sergio, Alecsander W. Zareb, Raizel M. Frasier, Sarah E. Wean, Phillip A. Starski, Angela Grippo, Frederic W. Hopf

## Abstract

Alcohol Use Disorder (AUD) remains a substantial global challenge, with notable individual and sex differences. Heart rate variability (HRV) offers a potential link into the autonomic system underlying alcohol drinking. While simpler, time-domain HRV metrics have been explored, non-linear indicators could provide unique insights into more complex states like compulsive drinking, intake that persists despite negative consequences. We examined heart rate traces from male and female rats during intake of alcohol-only and alcohol+10mg/L quinine, a putative compulsive drinking model. In particular, we re-analyzed previously published findings, but newly focused on several non-linear HRV metrics: the Poincare ratio (possibly reflecting sympathovagal balance) and MultiScale entropy (MSE, perhaps reflecting unpredictability in broader and narrower time windows), along with the previously unexamined spectral HRV metrics (the parasympathetic indictor high-frequency HRV, HF-HRV%). HRV was assessed for the first ∼1100 heart beats (∼2.5 minutes), to assess HRV correlates of greater alcohol front-loading while having approximately similar amount of data per session. Lower Poincare ratio and higher HF-HRV% were overall related to higher alcohol consumption. In contrast, compulsive drinking, but not alcohol-only, was associated with lower broader-window HRV entropy, which was more pronounced in males. Drinking- and sex-related patterns were further identified using Principle Component Analysis, Ordinary Least Squares, and Linear Mixed Models. Limitations included perhaps being underpowered to detect significant relationships, especially in females. Thus, our preliminary findings suggest that combining HRV metrics could represent useful biomarkers for consumption levels during front-loading, which distinguished general intake from compulsion.

## INTRODUCTION

Alcohol intake is now a leading cause of death in U.S. adults [1], and continues to be a significant global problem [2–4]. In addition, there are rising consumption rates [2, 5] and often greater alcohol-related problems among women [6, 7], highlighting the critical need to better understand sex [8] and individual [9, 10] differences in alcohol mechanisms when seeking to personalize and improve treatment. Further, compulsion for alcohol, where intake persists despite negative consequences, is one factor that can significantly contribute to problem drinking [11–13]. Thus, physiological biomarkers that predict alcohol intake could help develop more effective therapies [14–16]. We also speculate that compulsion models may provide a valuable, more general window into conflicted decision-making.

Heart rate variability (HRV) patterns are of wide interest as autonomic/arousal indicators, since these could help identify individual differences in arousal mechanisms and provide such biomarkers to perhaps improve treatment [14–16]. HRV is simple to measure in humans, and some metrics more clearly reflect sympathetic (SNS) influence (“fight or flight”) and others parasympathetic (PNS) (“rest and digest”) [17, 18]. A priori, several studies have noted that female mammals often have higher overall PNS influence, and males greater SNS relative influence, at rest and during some tasks (reviewed in [19]). For example, we previously showed that greater PNS HRV influence, indicated by rMSSD (root mean squared of the standard deviation), relates to higher alcohol intake and compulsion in female rats [20], while higher rMSSD relates to greater alcohol drives in female humans [21]. In contrast, male rat alcohol intake related to more basic SNS HRV markers [20], and there can be sex differences in human SNS stress responses [22]. However, the significance of sex differences and similarities in arousal patterns for alcohol and other behaviors remains unclear.

Importantly, several types of non-linear HRV indicators, like entropy, can often add additional and unique information [23–25]. Thus, the present data reflects a more thorough and deeper re-analyses of heart rate results that were first assessed in Frasier and colleagues [20]. In particular, the previous work focused on simpler “time” HRV metrics, while this new re-analysis focused on non-linear and spectral HRV metrics that might distinguish alcohol-only and compulsive drinking, as well as possible sex similarities and differences. Two non-linear HRV metrics (Poincare plot ratio and entropy), together with related spectral PNS measures (relative incidence of high-frequency HRV, HF-HRV%), distinguished alcohol-only from alcohol-quinine drinking, which has been considered to mirror some aspects of human alcohol compulsion [26, 27]. However, sample size and some other considerations limit interpretations from these findings, which should be considered preliminary and hypothesis generating rather than confirmatory, including where we cannot distinguish whether HRV-intake patterns are a consequence of longer-term drinking, or other impacts. Even so, these analyses provided additional foundation for use of HRV biomarkers to enhance human alcohol treatment, including incorporation of non-linear HRV metrics, which could open new insights into autonomic patterns and key underlying mechanisms that promote pathological motivation for alcohol and other conditions.

## METHODS

### Subjects and drinking methods

Animal care followed NIH guidelines and were approved by the Indiana University IACUC. Wistar rats arrived ∼PN50 and were single-housed in clear plastic cages. Similar number of males and females were in housing areas, under 12-h light/dark (dark 11:30 am-11:30 pm), with food and water *ad libitum*. Alcohol drinking began 2wks later, with all behaviors in dark phase.

As before [28, 29], rats first drank for ∼3mo under two-bottle-choice (20% alcohol v/v or water), with 16-24hr intake starting every Monday, Wednesday, and Friday at 1-2hr after lights out. Afterwards, rats switched to drink 20min/d, 5d/wk for ∼2mo, reaching ∼7-9mo old during studies. Then, in some sessions, rats drank alcohol-quinine (10mg/L), a putative compulsive drinking model [28, 29] (below). Specifically, all rats had 2-3 days drinking of alcohol-quinine to reduce novelty before testing, then drank alcohol-quinine ∼2 days/week and alcohol-only for other drinking days across testing. Alcohol-quinine intake was slightly (∼20%) reduced overall, compared to alcohol-only (t_(97)_=2.477, p=0.0150), as often observed. Finally, to measure HRV, rats were implanted with Stellar telemetry devices as described in [30], with 3 days recovery before recording HRV during alcohol drinking.

The present data reflects a more thorough and deeper re-analyses of heart rate traces during alcohol drinking, with a particular focus on the relation of consumption level and non-linear and spectral HRV metrics; previous work compared intake to rMSSD and other simpler HRV measures [20]. In addition, we present the results for each HRV metric in three ways. First, we had collected 1-3 drinking sessions per condition (alcohol-only or compulsive) per rat, and here we analyzed HRV patterns from all sessions, separately from animal identity. We [28, 29, 31, 32] and others [33–36] have done this before, including where we previously examined potential individual differences in lickometry [28] and HRV patterns [30] during alcohol drinking, and found variability in response pattern across rats, but little evidence for unique response patterns within particular rats. Even so, it is critical to understand whether any observed drinking-HRV patterns persisted when drinking sessions were averaged for each rat. Thus, we second collapsed drinking sessions for each rat, separately for alcohol-only and alcohol-quinine in each rat, and compared each HRV metric to drinking levels. Largely similar intake-HRV relations were observed. Thus, in this way, we were able to show which findings were most reliable, and, at the same time, retain a larger sample size of data for overall Principal Component Analysis (PCA, below). Third, we performed a Linear Mixed Effects model to retain within-rat results while also considering rat identity (Fig.S7).

Another important consideration is the description of quinine-resistant alcohol intake as compulsive. Quinine-resistant intake has been considered to mirror some aspects of human compulsion for alcohol [26, 27]. Also, while adulterating alcohol with quinine is used for convenience, similar salience network circuits mediate quinine- and shock-resistant alcohol drinking in rats [37], and several human studies link this circuit to alcohol compulsion [38–40], evidence in favor of quinine-resistance’s relevance to some aspects of human alcohol compulsion. In addition, we used 10mg/L quinine in alcohol, which we consider a moderate challenge, because it causes a relatively small drop in drinking relative to alcohol-only. In this way, we can determine different behavioral, autonomic, and brain mechanisms of compulsion versus alcohol-only with less confound from larger decreases in drinking levels with stronger negative consequences. Indeed, compulsion in humans is a clinical challenge because people maintain drinking despite the adverse consequences. In addition, using lickometry, we found that many different lick measures are significantly less variable during male moderate compulsive relative to alcohol-only [28]. We describe this as more stereotyped, automatic responding under compulsive conditions [41], consistent with automatic responding relating to human problem drinking [12, 42–45]. Interestingly, female rats show less variable licking, but only during the estrus stage [29], consistent with females overall using different response strategies relative to males [46].

Also important is that we did not examine blood alcohol levels, but previous work suggests that rats achieve 40-60mg/L blood alcohol [29, 47]. In addition, the long-term intermittent-access model we use exhibits several features considered to mirror human Alcohol Use Disorder, including escalating intake [12, 48], sensitivity to drugs that reduce human drinking [48–51], front loading (indicating motivation for alcohol) [28, 52, 53], and compulsive responding.

### Rationale for focusing on front-loading and a longer analysis window

Our primary objective was to determine HRV metrics across front-loading [28, 29], indicating stronger motivation to drink [52, 53], and more rapid alcohol intake in humans is associated with greater alcohol harm [54–57]. However, while rats typically drink most alcohol within 3-5 minutes, comparing HRV patterns involves analysis of HRV levels for similar duration across all sessions. Further, analysis of nonlinear measures like HRV entropy is more robust with a similar number of heart beats in each session. Several studies also emphasize that longer heart rate recordings are necessary to accurately determine non-linear HRV measures [58, 59]. Thus, 1100 Inter-Beat Intervals in each session was deemed a good compromise between capturing the majority of the frontloading period across all sessions, while having more reliable HRV analysis. In addition, the rate of drinking for alcohol-only and compulsive intake is similar [29, 32], indicating that similar time windows can be used for analyses comparing the two drinking conditions. Finally, while the total drinking session was 10 minutes [20], additional factors may contribute after front-loading including direct pharmacological effects of alcohol [60] and reduced motivation for further intake, which we sought to avoid.

### General HRV data analysis

Heartbeats were first converted to a train of Inter-Beat Times (IBTs, also called meanRR), the time between successive heartbeat R peaks; HRV reflects different aspects of variability in these IBTs [61]. Raw data underwent visual inspection and manual artifact removal, and sessions were only used if they had less than 2% of beats removed due to noise, as suggested [62]. [Fig.S1 shows a preliminary analysis of the impact of cutting out small bits of data on HRV metrics, with mostly little or only small effects.] HRV metrics were then calculated using Kubios (v.4.1, Kuopio, Finland) [18, 63, 64], with removal of non-stationary baseline shifts [65] and some automatic artifact correction (“medium” in Kubios) via cubic spline interpolation [66].

#### Specific HRV metrics

*The Poincare Plot ratio* is a non-linear geometric analysis. Each IBT was plotted against the subsequent interval (IBTₙ vs. IBTₙ₊₁) [24, 67] (**Fig.1A**). The cloud of points formed an ellipse, from which two standard deviations (SDs) were derived: SD1 (short-term variability) and SD2 (long-term variability) (**Fig.1A**). SD1 is similar to rMSSD, a well-validated index of PNS influence [17, 68, 69]. SD2 is considered to reflect contributions from both SNS and PNS activity, akin to the SD of IBTs (SDNN). SD2/SD1 is equivalent to SDNN/rMSSD and related to LF/HF (Low-Frequency vs High-Frequency HRV). Thus, SD2/SD1 can be considered a non-linear index of sympathovagal balance, with higher SD2/SD1 indicating greater SNS relative influence, e.g. where inhibiting SNS-promoting β-adrenergic receptors reduces Poincare ratio [70–72]. However, ratio can be influenced by factors such as cardiovascular health, age, and exercise [17, 23, 73] (although basal HRV is similar in alcohol-naïve and drinking rats [30]). Thus, caution is important when interpreting the meaning of HRV ratio or other metrics. Even so, positive affect and relaxation are linked to lower ratio and higher basic PNS indicators (like HF-HRV) in humans [74–78], and lower ratio relates to relief from restraint stress in rats [79]. Conversely, pain and stress can elevate ratio and decrease basic PNS measures [67, 80, 81]. Thus, the HRV ratio has the potential to indicate relative SNS-PNS balance under some conditions.

**Fig. 1.**
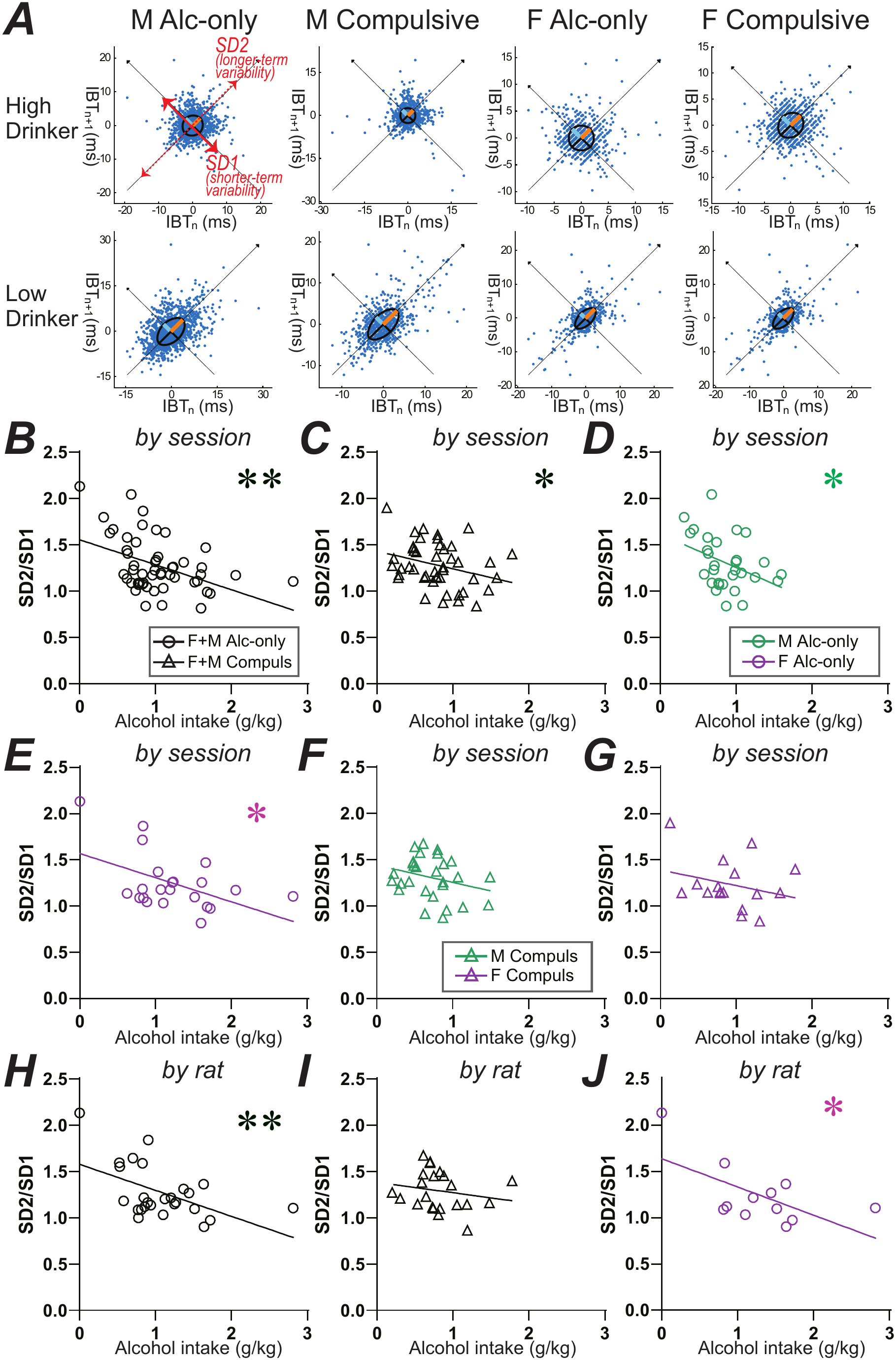
Greater alcohol drinking relates to reduced Poincare ratio. Male data in green, female in purple, black for sexes combined. (**A**) Examples of Poincare ratios from higher and lower drinkers from the four conditions (alcohol-only or compulsive, female or male). (**B,C**) Including all sessions, correlation of HRV ratio with (**B**) alcohol-only drinking (open circles) or (**C**) compulsive intake (triangles, alcohol with 10mg/L quinine). (**D-G**) Ratio and alcohol-only drinking in males (**D**) and females (**E**), and compulsive intake in males (**F**) and females (**G**). (**H-J**) Collapsed by rat, with males and females together, for alcohol-only (**H**) or compulsion (**I**), and (**J**) for female alcohol-only. F: female; M: male. *, ** for p<0.05, p<0.01 uncorrected.

*Spectral HRV metrics* have been widely utilized, using Fourier Transform analysis of different frequencies of IBT variability, as we have done before [72]. The power spectrum in rats is HF-HRV (0.75-2.5Hz), LF-HRV (0.19-0.75Hz), and VLF-HRV (0.02-0.19Hz). The raw power in each band (ms^2^) is then normalized to the total power (VLF+LF+HF), giving VLF-HRV%, LF-HRV% and HF-HRV% [80]; here, we only examined HF-HRV% as an indicator of PNS influence, since β-adrenergic receptor antagonism significantly increases HF-HRV% [72, 82–84].

*Entropy* measures assess randomness in data sets, where lower entropy reflects better predictability based on the recent past, and higher entropy indicating greater randomness (**Fig.3A**) [23, 24, 85]. We examined MultiScale entropy (MSE) (**Fig.3B-F**), which is built from the Sample entropy: with “a sample of length N, the SampEn is defined as the negative natural logarithm of the probability that if two sets of simultaneous data points of length m have distance < r, then two sets of simultaneous data points of length m + 1 also have distance < r;” with the scaling parameters m (embedding dimension; using default value 2 beats) and r (tolerance value; default: using 0.2×SD adjustment in relation to standard deviation of RR data) (Kubios, [24, 86]). MSE then “coarse-grains” the data (**Fig.3B**; see Fig.2 of [86] and Section 2.3 of [87]): MSE1 is from the original set of IBTs, while MSE2 involves averaging each pair of IBTs into one IBT, across the session; MSE3 involves averaging each three IBTs into one IBT; etc (**Fig.3B**); MSE reduces variability (by averaging) and decreases sample size. Sample entropy is then determined for each MSE level. Also, for a reliable estimate of Sample Entropy, at least 10^m^ data is needed (although, more conservatively, even 30 ^m^) [24]. Thus, in MSE analysis, this requires 100 (10 ^2^) for the sample entropy at each coarse-grained level. With 1100 data points, MSE11 will be coarse-grained to 100 data points. Importantly, these smaller data sizes would increase uncertainty in estimating broader-window entropy, but this influence would likely act similarly for compulsion and alcohol-only. Instead, we observed striking differences in broader entropy’s relation to drinking, especially reduced compulsive intake with higher entropy but no entropy relation to alcohol-only (**Fig.4**). Thus, uncertainty in broader-window estimates was not likely responsible for the broader-window entropy anti-compulsion relation, especially in males. In addition, however, data by rat were only trend significant for male compulsion relating to lower broader entropy (*p*=0.0532, **Fig.4I**). Thus, while our findings likely capture important aspects of arousal physiology in relation to different types of alcohol drinking, they should be considered preliminary and hypothesis-generating rather than confirmatory and conclusive.

**Fig. 2.**
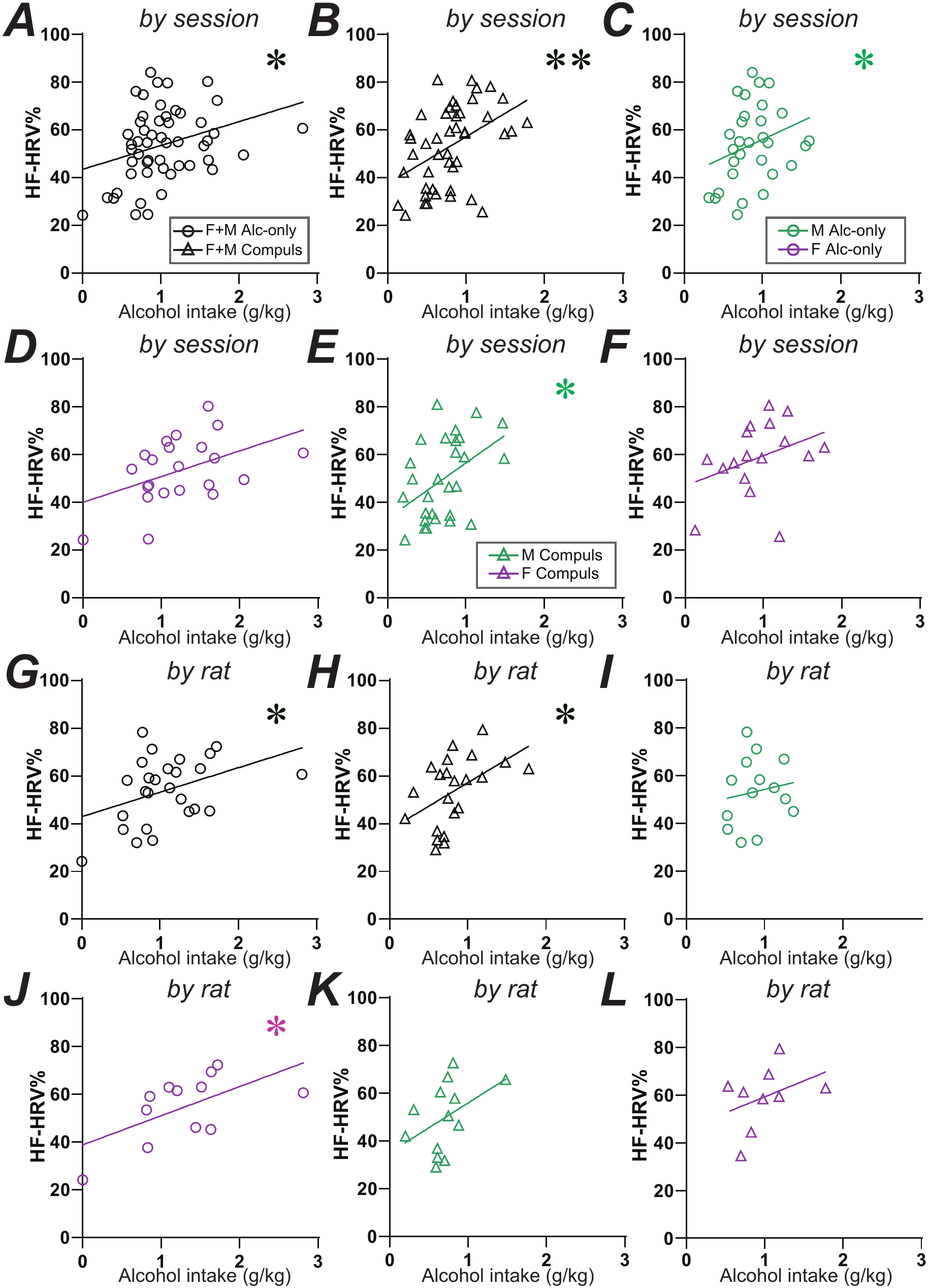
Greater alcohol drinking relates to higher HF-HRV%. (**A,B**) Comparison of drinking level and HF-HRV%, with male and female sessions combined, for (**A**) alcohol-only and (**B**) compulsive drinking. (**C-F**) Session data separated by sex for (**C**) male and (**D**) female alcohol-only, and (**E**) male and (**F**) female compulsion. (**G-L**) Sessions collapsed by rat. (**G,H**) Females and males together for (**G**) alcohol-only and (**H**) compulsion. (**I-L**) By rat, alcohol-only drinking in males (**I**) and females (**J**), and compulsive intake in males (**K**) and females (**L**). F: female; M: male. *, ** for p<0.05, p<0.01 uncorrected.

**Fig. 3.**
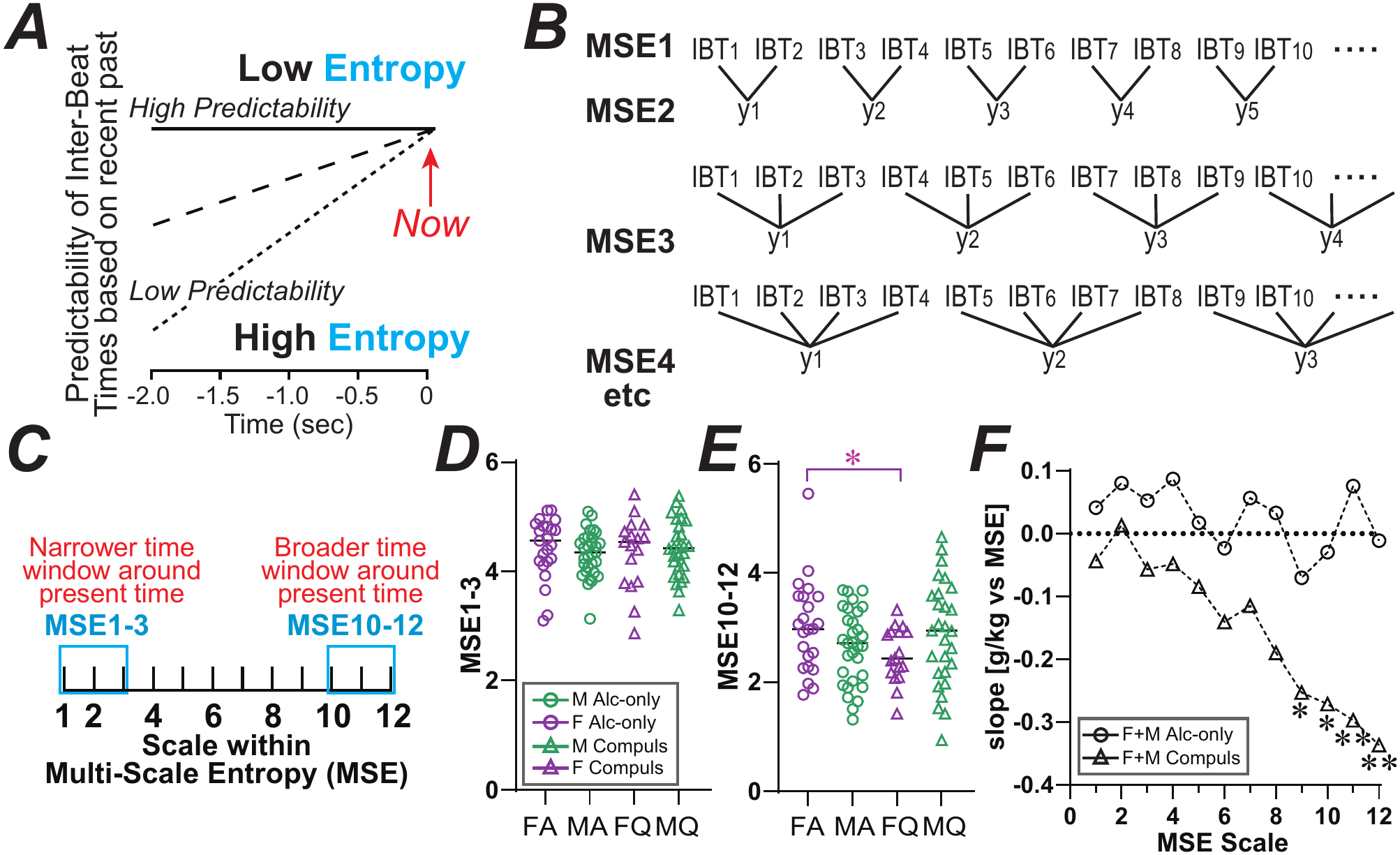
Measurement of different types of HRV entropy in relation to drinking. **(A**) Entropy measures indicate how well a given data point can be predicted from recent past results; thus, they assess randomness or chaos within a data set. (**B**) Cartoon of “coarse-graining” of Inter-Beat Times for Multi-Scale Entropy MSE1 through MSE4. For each successive MSE scale, a broader set of the original Inter-Beat Times is averaged. Entropy is then calculated from the resulting data series. (**C**) MSE assesses entropy across different time scales, here examining narrower time scales (MSE1-3) or broader time scales (MSE10-12), see Methods. (**D,E**) Average values of different MSEs did not differ across (**D**) male alcohol-only vs compulsion, but (**E**) did for females. (**F**) shows slope relation between a given MSE and alcohol intake, combining males and females, and separately for alcohol-only (circles) and compulsion (triangles). F: female; M: male. *, ** for p<0.05, p<0.01 uncorrected.

**Fig. 4.**
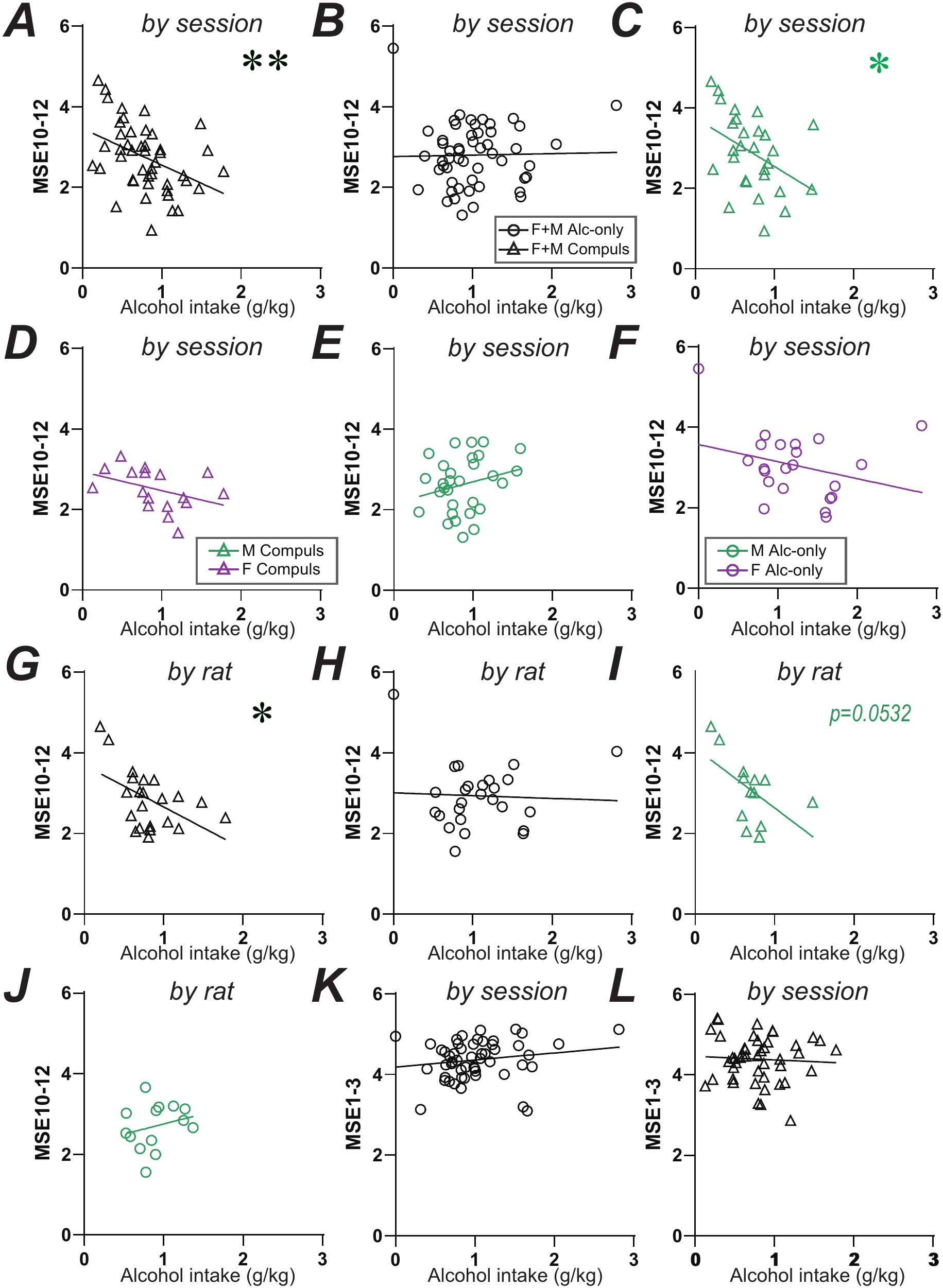
Relation of HRV entropy metrics to alcohol drinking. (**A,B**) Analyzed by session and combining males and females, broader-window MSE (entropy in a wider time window around the present) in relation to (**A**) compulsive and (**B**) alcohol-only intake. (**C-F**) The compulsion-MSE10-12 relation was driven by (**C**) males rather than (**D**) females, and with no relation to alcohol-only in (**E**) males or (**F**) females. (**G-J**) Analyzed by rat with sexes combined, broader-window entropy relation to (**G**) compulsion but not (**H**) alcohol-only, with a similar pattern in males only for (**I**) compulsion versus (**J**) alcohol-only. (**K,L**) Analyzing by session and combining sexes, narrow-window entropy (MSE1-3) did not relate to (**K**) alcohol-only or (**L**) compulsion. F: female; M: male. *, ** for p<0.05, p<0.01 uncorrected.

In addition, our choice of MSE1-3 and MSE10-12, rather than other MSE windows, was to provide some measure of narrower- and broader-window HRV entropy, but otherwise was somewhat arbitrary. Previous studies have used a variety of MSE windows to examine different aspects of entropy, e.g. [88] used MSE1-5 for shorter and MSE6-10 for longer window entropy (although this is from human, with slower heart rate than rat). However, using somewhat different MSE windows led to similar results as MSE10-12 and MSE1-3 (not shown).

Importantly, HRV entropy largely reflects PNS influences outside of cardiovagal inputs, for example misfiring in heart muscles [89]. Evidence of this comes from use of the cardiovagal inhibitor atropine, which strongly reduces HF-HRV at doses which have only limited effects on HRV entropy [90–92]. In agreement, HRV entropy can be in a different principal component from cardiovagal HRV frequency metrics [23].

### Data Analysis and Statistics

MeanRR was non-normal and thus log-transformed, while ratio, HF-HRV%, and MSE metrics were not log transformed; ratio was normally distributed for all groups except female alcohol-only. Simpler statistics were analyzed using GraphPad Prism. Our primary statistical comparison was correlation with alcohol drinking level, which was compared across all sessions, separately for alcohol-only versus alcohol-quinine (with males and females combined), and also separately by sex and condition (male alcohol-only, male compulsive, female alcohol-only, female compulsive). Females drank more alcohol (females: 1.09±0.08g/kg; males: 0.79±0.04g/kg; t_(97)_=3.501, *p*=0.0007), as often observed [8].

Also, we did not perform corrections for multiple comparisons, and instead report the raw p values. The PCA and ordinary least-squares (OLS) analyses (below) then helped clarify which of the overall HRV patterns most clearly correlated with drinking levels. Finally, as detailed in Fig.S2, there were no sex differences in the levels of different HRV metrics except for smaller meanRR in females overall (the widely observed faster heart rate in females), and MSE10-12 in female alcohol-only vs compulsion (below).

*PCA* was applied across HRV measures and heart rate. Variables were first z-scored (standardized to mean=0, SD=1). Component loadings are reported as correlation loadings. The number of components retained was guided by scree plot, with PC1 and PC2 giving cumulative variance of ≥ 80% (Fig.S3). First, a single PCA was performed across all drinking sessions to characterize the global covariance structure of the HRV measures. and to generate individual PC scores for each animal. One-way ANOVA assessed group-level differences by sex and drinking condition for PC1 and PC2 scores.

We then examined the relationship between PC scores and alcohol drinking level separately for each group, to determine whether individual HRV metrics contributed to each PC differently for sex and/or drinking condition. PC-intake associations were assessed in two complementary ways: (1) bivariate Pearson correlations computed separately within each group, and (2) OLS multiple regression with PC1 and PC2 entered simultaneously as predictors of alcohol intake, estimated separately per group. OLS examines the contribution of each PC while accounting for the influence of the other PC. PCA, OLS, and LME were performed in Python using scikit-learn (v1.x; PCA and standardization), NumPy (OLS), statsmodels MixedLM (LME), and SciPy (statistical testing), with statistical significance set at α=0.05. Fig.S4 shows a leave-one-out analysis suggesting differing impacts of the varied HRV metrics on drinking across sex and drinking condition.

## RESULTS

### Overall, higher alcohol drinking related to lower HRV Poincare ratio and greater HF-HRV%

We first looked at the HRV Poincare ratio, which may be most related to sympathovagal balance [17, 71]. In addition, previous work in humans has linked PNS more to female responding and SNS more to male responding (reviewed in [19]), and for alcohol in rats [20]. Thus, we hypothesized that higher male drinking would correlate with higher ratio, and greater female intake with lower ratio. Instead, lower HRV ratio correlated with higher drinking levels overall (**Fig.1**). Similarly, we examined the spectral PNS HRV metric, HF-HRV%, with complementary a priori hypotheses as ratio (drinking related to greater HF-HRV in females and lower in males), and also found some unexpected findings (**Fig.2**). In addition, as detailed in Methods, we present HRV-intake correlations in two ways, one considering all drinking sessions separately (“by session” in figures) and one collapsing different sessions from each rat into a single value each for alcohol-only and compulsive intake (“by rat” in figures).

Across all sessions, lower Poincare ratio SD2/SD1 strongly predicted greater alcohol consumption (F_(1,95)_=15.50, *p*=0.0002). **Fig.1A** shows examples of Poincare ratios separated by high or low drinking, sex, and alcohol-only versus compulsion. Higher drinkers had a rounder Poincare distribution (larger SD1 and smaller SD2), while lower drinkers had a more ellipsoid distribution (larger SD2 and smaller SD1). SD1 has been linked more to PNS, and SD2 to higher SNS influence, while inhibiting SNS-promoting βARs shifts the distribution from ellipsoid to round suggesting greater PNS [70–72]. Thus, under some conditions, greater Poincare ratio may reflect higher SNS influence (see Methods).

Across sessions with sexes combined, greater drinking was significantly correlated with lower ratio for alcohol-only (**Fig.1B**; F_(1,50)_=11.84, *p=*0.0012) and compulsive intake (**Fig.1C**; F_(1,43)_=4.21, *p=*0.0463). Separated by sex, lower ratio correlated with greater alcohol-only drinking in males (**Fig.1D**; F_(1,28)_=5.064, *p=*0.0325) and females (**Fig.1E**; F_(1,20)_=6.411, *p=*0.0198) and but did not relate to compulsive intake in either sex alone (**Fig.1F,G**; male: F_(1,26)_=2.057, *p=*0.1634; female: F_(1,15)_=1.193, *p=*0.2920). When examined by rat with sexes collapsed, lower HRV ratio correlated with higher intake overall (F_(1,46)_=9.741, *p*=0.0031) and to lower alcohol-intake (**Fig.1H**; F_(1,24)_=9.02, *p=*0.0062) but not compulsive drinking (**Fig.1I**; F_(1,20)_=0.7355, *p=*0.4013). Further, with the smaller sample size and power, lower ratio was related to higher intake for female alcohol-only (**Fig.1J**; F_(1,10)_=6.306, *p=*0.0309) but not male alcohol-only (Fig.S5A; F_(1,12)_=1.369, *p=*0.2647) or compulsion in males (Fig.S5B; F_(1,11)_=0.4208, *p=*0.5298) or females (Fig.S5C; F_(1,7)_=0.03442, *p=*0.8581). Taken together, these results suggest the possibility that higher HRV ratio was related to lower alcohol drinking for alcohol-only drinking in both sexes, with less clear impact for compulsive intake.

Several drinking-HRV relationships were also observed when examining the spectral HRV metric HF-HRV%, widely linked to PNS impacts [17, 68, 69], and predicted to be the converse pattern of ratio. Greater drinking related to higher HF-HRV% across sessions with sexes combined (F_(1,95)_=14.01, *p=*0.0003). However, compared to HRV ratio, higher HF-HRV% was related to greater alcohol-only intake (**Fig.2A**; F_(1,50)_=5.401, *p=*0.0242), and even more so to compulsive drinking (**Fig.2B**; F_(1,43)_=10.81, *p=*0.0020). Separated by sex, higher HF-HRV correlated with greater alcohol-only drinking in males (**Fig.2C**; F_(1,20)_=5.186, *p=*0.0339) but not females (**Fig.2D**; F_(1,28)_=2.51, *p=*0.1244), and to compulsive intake in males (**Fig.2E**; F_(1,26)_=6.721, *p=*0.0154) but not females (**Fig.2F**; F_(1,15)_=2.172, *p=*0.1612). When examined by rat, collapsed by sex, greater HF-HRV% was correlated with higher alcohol drinking levels overall (F_(1,46)_=9.176, *p*=0.0040), and to greater alcohol-only (**Fig.2G**; F_(1,24)_=4.502, *p=*0.0444) and compulsive drinking (**Fig.2H**; F_(1,20)_=6.207, *p=*0.0216). Further, higher HF-HRV% was not related to higher alcohol-intake intake in males (**Fig.2I**; F_(1,12)_=0.2834, *p=*0.6042), but did correlate with female alcohol-only (**Fig.2J**; F_(1,10)_=5.542, *p=*0.0404), although not compulsion in males (**Fig. 2K**; F_(1,11)_=2.567, *p=*0.1374) or females (**Fig.2L**; F_(1,7)_=1.183, *p=*0.3128) but with trends in both. Together, these results support the hypothesis that higher PNS, expressed as greater HF-HRV%, was related to higher alcohol drinking, especially compulsion.

### Compulsive but not alcohol-only drinking relates to some aspects of HRV entropy

We next examined several HRV entropy measures (which are not log transformed). More generally, entropy metrics assess randomness within a data set, especially how well a given data point can be predicted from recent past results (**Fig.3A**) [25, 85]. In particular, we examined MultiScale entropy (MSE). As detailed in Methods, this involves “coarse-graining” heartbeat values to examine entropy across narrower versus broader windows of time. Specifically, the original train of Inter-Beat Times (“MSE1” in **Fig.3B**) was averaged for every two Inter-Beat Times (MSE2), every three Inter-Beat Times (MSE3), etc, and entropy was then calculated for each coarse-grained scale. One way to conceptualize MSE results is that MSE1-3 examines entropy over narrower time scales, where MSE1 is closest to real time (**Fig.3B,C**). Alternately, MSE10-12 examines entropy across a much broader time window (**Fig.3C**), and we term MSE10-12 and MSE1-3 as “broader-window” and “narrower window” entropy. In addition, MSE values showed little difference across groups (**Fig.3D,E**), although females showed reduced broader-window MSE during compulsion vs alcohol-only (**Fig.3E**; t_(38)_=2.226, *p*=0.0320). Instead, as shown below, entropy measures correlated with drinking level differently based on drinking condition (challenged or not), even with few average differences.

A priori, we hypothesized that broader-window MSE HRV would relate to reduced compulsive drinking, with little relation to alcohol-only. In particular, we have speculated that compulsive intake involves removal of attention from negative consequences in order to maintain focus on the associated reward [26, 28]. In contrast, broader-window entropy can associate with challenge conditions, including slower reaction time in harder tasks [86, 93], which could reflect evaluation [94] or awareness [88]. Though highly speculative, in this framing, higher awareness of actual contingencies (perhaps indexed in broader-window entropy) might reduce the ability to maintain compulsive intake.

As predicted, higher broader-window entropy (MSE10-12) correlated with significantly lower compulsive drinking but not alcohol-only, with no overall relation to intake across all sessions combined (F_(1,95)_=2.085, *p*=0.1521). Across sessions, collapsed across males and females, broader-window entropy was correlated with lower compulsive drinking (**Fig.4A**; F_(1,43)_=8.977, *p*=0.0045) but not alcohol-only intake levels (**Fig.4B**; F_(1,50)_=0.0239, *p*=0.8778), and with a significant difference in slopes between compulsion and alcohol-only (F_(1,93)_=6.065, *p*=0.0156). The compulsion-MSE10-12 relation was driven by males (**Fig.4C**; F_(1,26)_=5.436, *p=*0.0277; **Fig.4D**; females: F_(1,15)_=2.800, *p=*0.1150), with no relation of broader entropy for alcohol-only in males (**Fig.4E**; F_(1,28)_=1.707, *p=*0.2020) or females (**Fig.4F**; F_(1,15)_=2.800, *p=*0.115). In addition, there was a significant difference in the entropy-intake slope between alcohol-only and compulsive intake for males only (F_(1,54)_=6.996, *p*=0.0107) and for males and females combined (F_(1,93)_=6.065, *p*=0.0156), again suggesting that broader-window entropy related differently to compulsion and alcohol-only. **Fig.3F** shows the slope relationship and significance for each MSE level individually.

A similar pattern was observed by rat, where higher broader-window entropy related to lower compulsive drinking (**Fig.4G**; F_(1,20)_=6.974, *p=*0.0157) but not alcohol-only (**Fig.4H**; F_(1,24)_=0.05206, *p=*0.8215), with a trend for differences in slopes (F_(1,44)_=3.322, *p*=0.0752). While this tended to be driven by male compulsion, this was only trend significant in males (F_(1,11)_=4.691, *p*=0.0532). On the other hand, male broader-window entropy did not relate to alcohol-only (F_(1,12)_=0.760, *p*=0.4004), but there was a difference in slope between male compulsion and male alcohol-only (F_(1,23)_=4.902, *p*=0.0370); there was also no broader-window entropy relation for female alcohol-only or compulsion by rat (Fig.S5D,E). Together, these indicate that broader-window HRV entropy was negatively associated with compulsive drinking levels, with no relation for alcohol-only. Also, this might reflect more prominent effects in males, although the lower sample size in females precludes definitive statements about non-effects in female compulsion under these conditions.

To better understand the broader-window entropy findings, we compared results with narrower-window HRV entropy (MSE1-3), which has been related more to cognitive flexibility (see Discussion). Narrower-window entropy did not relate to alcohol-only (**Fig.4K**; F_(1,50)_=1.499, *p=*0.2265), and, unlike broader-window entropy, did not relate to compulsive intake (**Fig.4L**; F_(1,43)_=0.1435, *p=*0.7067) (data by rat and sex shown in Fig.S6). Thus, broader-but not narrower-window entropy related to reduced compulsive drinking.

### Heart rate and alcohol intake

We also examined the relation between drinking and the log-transformed Inter-Beat Time (meanRR), the reciprocal of heart rate. Heart rate and HRV can relate differently to behavior and affective state [95, 96]. In sessions collapsed by sex, lower meanRR (more rapid heartbeats) was related to greater compulsive (**Fig.5A**; F_(1,43)_=7.810, *p*=0.0077) but not alcohol-only (**Fig.5B**; F_(1,50)_=1.671, *p*=0.2020), although with no slope difference between compulsion and alcohol-only (F_(1,93)_=2.556, *p*=0.1133). In addition, in male sessions, meanRR related to less compulsive drinking (**Fig.5C**; F_(1,26)_=5.153, p=0.0317) but not alcohol-only (**Fig.5D**; F_(1,28)_=0.028, p=0.8685), with no slope difference between compulsion and alcohol-only (F_(1,54)_=2.993, *p*=0.0894), and no clear significance for female alcohol-only (**Fig.5E**; F_(1,20)_=0.123, *p*=0.7298) or compulsion (**Fig.5F**; F_(1,15)_=0.691, *p*=0.4191). For by-rat collapsed by sex, lower meanRR was related to greater compulsive intake (**Fig.5G**; F_(1,20)_=7.771, *p*=0.0114) but not alcohol-only drinking (**Fig.5H**; F_(1,20)_=1.137, *p*=0.2969). Further, meanRR by rat was significantly related to male compulsive intake (**Fig.5I**; F_(1,11)_=9.575, *p*=0.0102), but not alcohol-only (**Fig.5J**; F_(1,12)_=0.077, *p*=0.7859), and with no clear significance for female alcohol-only (**Fig.5K**; F_(1,10)_=0.107, *p*=0.7508) or compulsion (**Fig.5L**; F_(1,7)_=0.051, *p*=0.8271). However, there was no significance for slope differences between compulsion and alcohol-only, collapsed by sex (F_(1,44)_=3.112, p=0.0847) and for males (F_(1,23)_=3.471, p=0.0753) by rat. These findings, across the first ∼2.5min of drinking, replicated where male compulsive intake correlated with meanRR over the whole 10 minute drinking session, as reported previously in this data set [20].

**Fig. 5.**
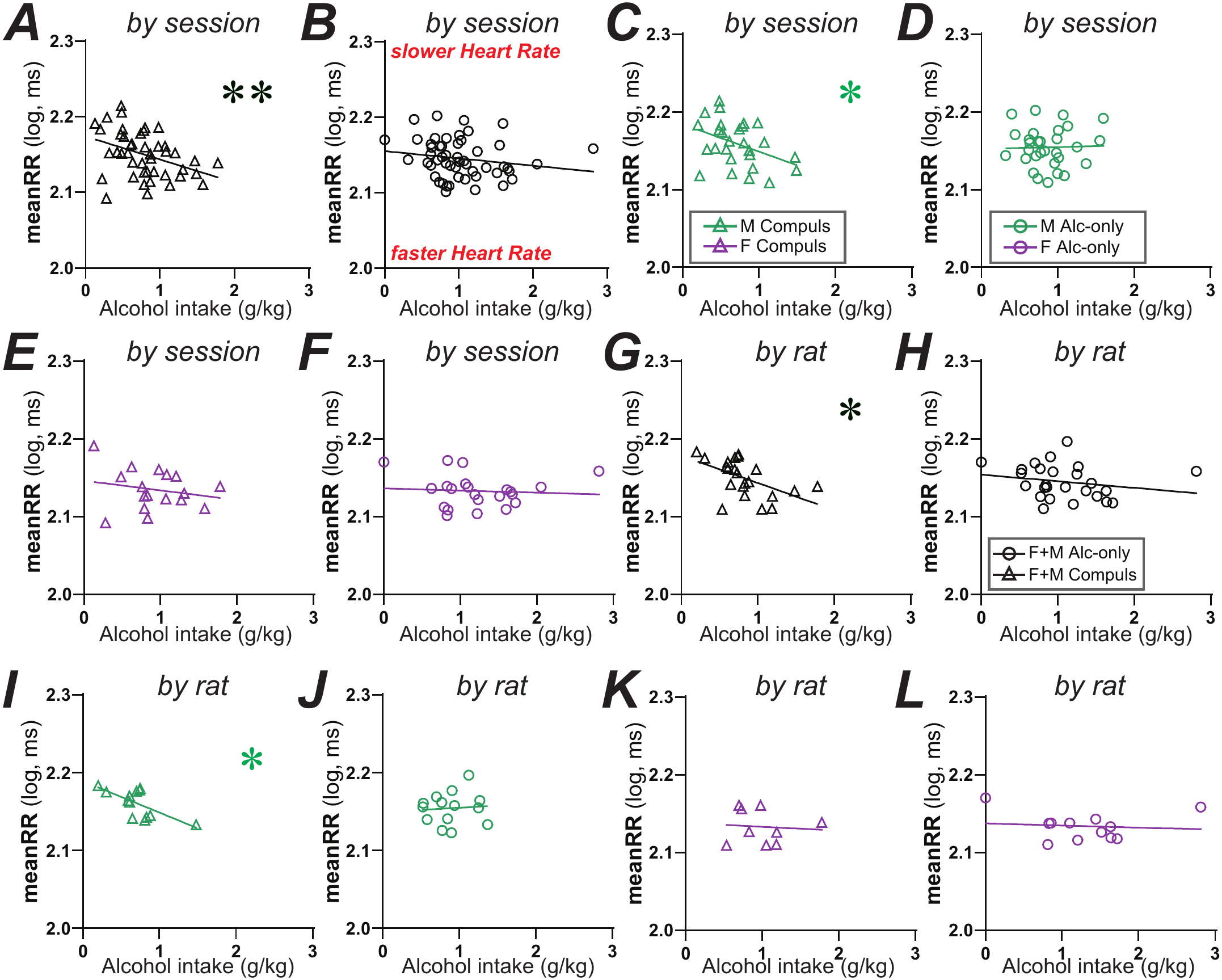
Time between heartbeats (meanRR) related to compulsion-like intake, especially in males. (**A,B**) By session, more time between heartbeats (larger meanRR) related to lower (**A**) compulsive but not (**B**) alcohol-only drinking. (**C-F**) Drinking level related to (**C**) male compulsion rather than (**D**) alcohol-only, or female (**E**) compulsion or (**F**) alcohol-only. (**G,H**) By rat collapsed by sex, more rapid heartbeat (lower meanRR) related to (**G**) more compulsive intake with (**H**) no relation to alcohol-only. (**I-L**) By rat, drinking level related to (**I**) male compulsion rather than (**J**) alcohol-only, or female (**K**) compulsion or (**L**) alcohol-only. F: female; M: male. *, ** for p<0.05, p<0.01 uncorrected.

### Principle Component Analysis validated the impact of different HRV metrics on drinking levels

We then applied PCA to characterize the variance among meanRR and the four HRV metrics in relation to drinking, using the results separated by session. Two PCs were retained based on the scree plot, together explaining ≥80% of variance (Fig.S3). PC1 explained 48.6% of overall variance and received negative loadings for ratio and meanRR and positive loading for HF-HRV% (**Fig.6A**). PC2 explained 32.9% and captured a positive relation with the two MSE metrics (**Fig.6A**). Also, one-way ANOVA revealed a significant effect of sex on PC1 scores (F_(1,95)_=4.19, *p*=0.0434), with higher values in females. PC2 had no sex effects (F_(1,95)_=0.01, *p*=0.9261), nor did drinking condition (alcohol-only vs. compulsive, collapsed across sex) for PC1 (F_(1,95)_=0.04, *p*=0.8398) or PC2 (F_(1,95)_=0.00, *p*=0.9690).

**Fig. 6.**
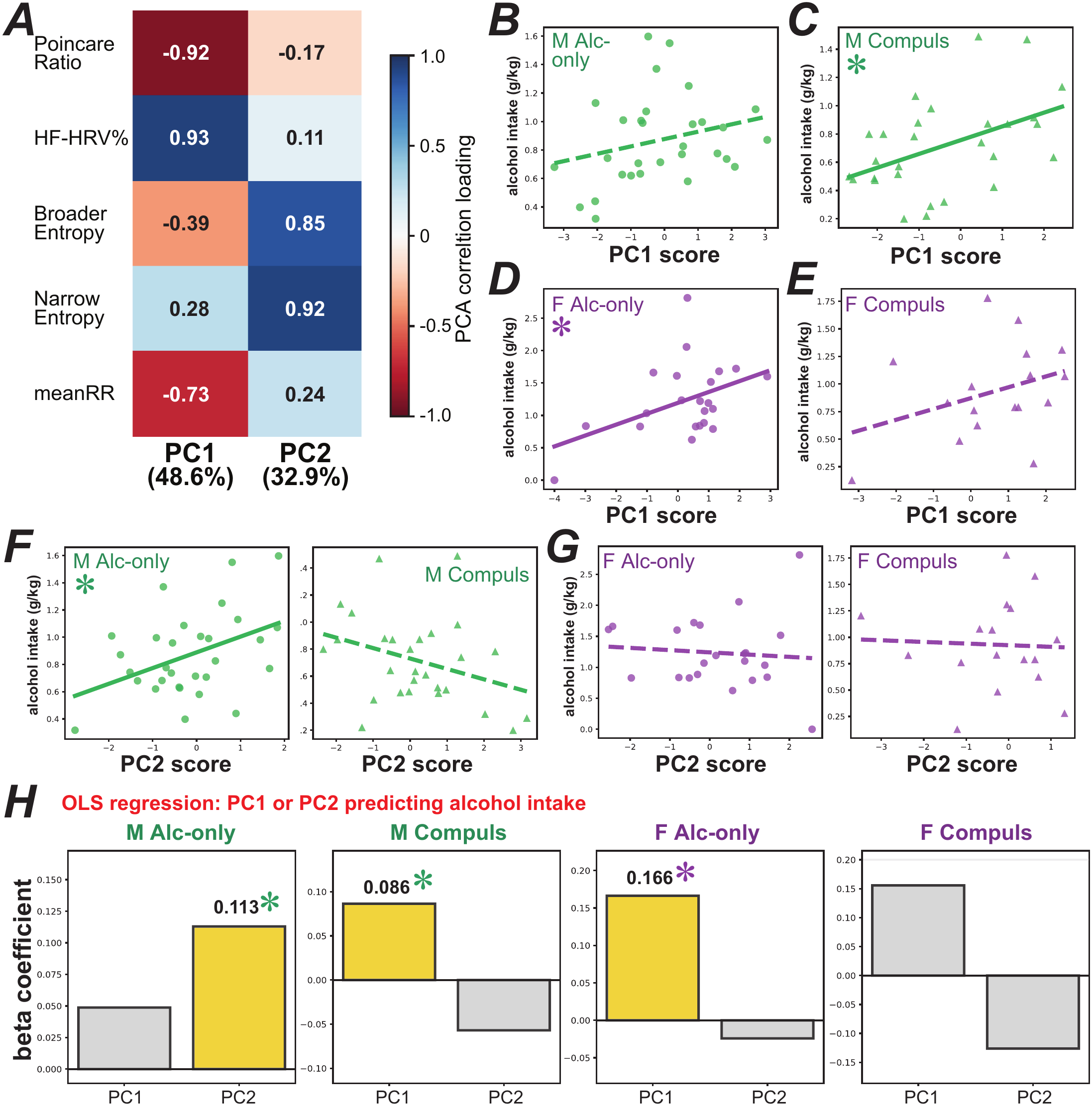
PCA and OLS validate how inclusion of non-linear and spectral metrics unmasks different HRV patterns related to drinking condition and sex. **(A)** PC loadings for each metric examined. **(B-E)** PC1 relation to male **(B)** alcohol-only and **(C)** compulsive intake, and to female **(D)** alcohol-only and **(E)** compulsive intake. **(F,G)** PC2 relation to **(F)** male and **(G)** female drinking**. (H)** Results from OLS with PC1 and PC2 as predictors. Yellow bars indicate significant beta coefficient. F: female; M: male. *, ** for p<0.05, p<0.01 uncorrected.

Pearson correlations between PC scores and alcohol intake were computed separately per group (**Fig.6B-G**). PC1 correlated positively with male compulsive drinking (**Fig.6C**; *r*=0.459, *p*=0.0141) but not alcohol-only intake (**Fig.6B**; *r*=0.264. *p*=0.1583), and also significantly related to female alcohol only (**Fig.6D**; *r*=0.439, *p*=0.0409) but not compulsion (**Fig.6E**; *r*=0.351, *p*=0.1678). These agree with observations above that higher drinking was related to lower ratio and higher HF-HRV% (**Fig.1**,**2**), and also that male compulsive intake was greater with faster heart rate (**Fig.5C**). Furthermore, PC2 was positively linked to the two entropy metrics, and correlated with male alcohol-only drinking (**Fig.6F**; *r*=0.421, *p*=0.0206) with a trend for negative correlation with male compulsion (**Fig.6F**; *r*=-0.331, *p*=0.0855), concurring with male compulsion relating to lower broader-window entropy (**Fig.4C**); narrow-window entropy also trend related to male alcohol-only (Fig.S6A). No significant PC2-drinking associations were observed for females (**Fig.6G**; alcohol-only: *r*=-0.087, *p*=0.7016; compulsion: *r*=-0.044, *p*=0.8668). Together, these PCA results validate that male alcohol intake was related to multiple HRV metrics, with fewer significant findings for females. In addition, using a leave-one-out PCA analysis, removing one of the HRV metrics had limited effects on the overall significance of the model (Fig.S4).

Ordinary Least Square (OLS) multiple regression with PC1 and PC2 as simultaneous predictors confirmed these patterns (**Fig.6H**). The model was significant for male PC1 for alcohol-only (adjusted *r*^2^=0.18, F=4.24, *p*=0.025) and compulsion (adj. *r*^2^=0.21, F=4.56, *p*=0.020), with PC1 the main predictor of male compulsion and PC2 for male alcohol-only. Overall, OLS was not significant in females for alcohol-only (adj. *r*^2^=0.11, F=2.32, *p*=0.126) or compulsion (adj. *r*^2^=0.10, F=1.86, *p*=0.192), There were also significant interactions from OLS, for PC1 in female alcohol-only (β=0.166, *p*=0.0482) and male compulsion (β=0.086, *p*=0.0288), and for PC2 in male alcohol-only (β=0.113, *p*=0.021). Taken together, these analyses validate that several non-linear plus spectral HRV metrics during alcohol drinking were significantly correlated with drinking levels, with differences between alcohol-only and compulsion as well as some sex differences.

### Linear Mixed Effect model analysis of alcohol-HRV patterns

Finally, we performed an LME analysis, where data points within-rat are considered along with subject identity (Fig.S7). Observed patterns were largely similar as other analyses. With sexes combined (Fig.S7A), lower ratio related to greater alcohol-only intake (*p*=0.0009) but less so for compulsion (*p*=0.0374), while higher HF-HRV% related to more alcohol-only (*p*=0.0048) and compulsion (*p*=0.0026). Broader-window entropy also related to lower compulsive intake (*p*=0.0015) but not alcohol-only (*p*=0.7403), with *p*=0.0015 for differences in slope between alcohol-only and compulsion (Fig.S7B) (and *p*=0.0211 for male compulsion). MeanRR (time between beats) also related to lower compulsive intake (*p*=0.0080) rather than alcohol-only (*p*=0.1195) (and *p*=0.0248 for male compulsion). Also, using PCA and OLS, PC1 and PC2 loadings (Fig.S7C,D) were similar to **Fig.6**, while (Fig.S7D-F) PC1 only related significantly to male compulsion (*p*=0.0062), with trends for other conditions, and PC2 only related to male alcohol-only (*p*=0.0160). Also, OLS analyses related to male alcohol-only and compulsion (Fig.S7G), with trends for females. Taken together, this LME analyses uncovers key aspects of alcohol-HRV relations mostly in line with as previous analyses.

## DISCUSSION

We provide new evidence that combining HRV metrics has the potential to provide separable candidate HRV signatures for alcohol-only versus compulsive drinking, and perhaps in males versus females. Alcohol addiction remains a substantial problem, and validated biomarkers like HRV metrics could help personalize and improve treatment. We and others have compared alcohol behaviors in rats and humans with simpler HRV measures [19–21], but non-linear indicators often add additional, unique information [23–25]. The nonlinear Poincare ratio and spectral HF-HRV% were the strongest overall predictors of drinking level, and suggested that greater PNS influence related to higher consumption in both sexes overall. In contrast, reduced entropy in a broader time window correlated with greater compulsive drinking but not alcohol-only intake, especially in males. PCA and OLS largely validated these analyses. One limitation is that we may be underpowered to detect significant relationships, especially in females. Even so, our studies suggest the possibility that incorporating non-linear with other HRV metrics might together provide useful indicators of drive for alcohol under different drinking conditions.

The strongest overall effects were in the HRV Poincare plot and HF-HRV%, where lower ratio and higher HF-HRV% (both putative indicators of greater PNS relative influence) were linked to greater alcohol consumption. Lower ratio is linked to relaxation and positive affect in humans [77, 78], and to digestion and relief from stress in rats [79, 97]. Also, lower HRV ratio correlates with greater male engagement in an anxiety-like behavior, suggesting reduced anxiety [30]. However, interpretation of HRV ratio (and HRV metrics more generally) must be done with caution, and interpreting ratio as SNS-PNS balance is complicated under conditions such as aging, exercise, and cardiovascular disease [17, 23, 73]. Notably, a train of heart beats reflects the integration of many different SNS and PNS influences. For example, strong exercise increases heart rate and decreases HRV, but it can be problematic to interpret HRV shifts as SNS versus PNS when faster heart rate has a dominant impact on the integrated heart beat train [17, 23, 73]. Nonetheless, even with these caveats, HRV can provide useful insights into arousal changes related to behavior under some conditions, and it will be interesting to relate alcohol intake levels and HRV ratio in human problem drinkers. Also, we note that Poincare ratio is equivalent to SDNN/rMSSD, which we previously examined for the first minute of drinking [20]; this ratio did not relate to alcohol drinking levels, but males and not females had a significant increase in ratio at drinking onset. Importantly, our findings together thus suggest that ratio values at drinking onset are less related to total intake, while greater PNS (lower ratio) across the front-loading period do predict higher consumption.

We note that HF-HRV% showed some comparable relationships to drinking level as HRV ratio, although some patterns diverged, with HRV ratio perhaps more related to alcohol-only intake (**Fig.1**) and HF-HRV% to compulsion (**Fig.2**). Indeed, while HRV ratio and HR-HRV% are clearly related, they likely index somewhat different aspects of arousal regulation. HRV ratio has been considered to reflect a sympathovagal balance, which has the potential to be more nuanced without larger changes in SNS or PNS per se. As such, ratio can relate better to anxiety than HF-HRV or LF-HRV [81], while prefrontal areas in humans regulate both SNS and PNS [88, 98, 99] and can link to ratio more than basic metrics [100]. This could also perhaps reflect where the Poincare HRV ratio compares adjacent Inter-Beat Times, versus HF-HRV which is derived from longer-term cyclicity of HRV levels, reflecting variability across several hundred milliseconds versus several seconds or more in rats.

HRV entropy was associated with different aspects of drinking than ratio and HF-HRV. In particular, higher broader-window HRV entropy was correlated with reduced compulsive drinking, possibly with a greater impact in males, but was not associated with alcohol-only intake. Narrower-window entropy showed less consistent relationships with intake level (Fig.S6). Previous human studies may provide some context. Sample entropy (MSE1) in humans is often elevated during mental challenges versus simpler tasks or rest [25, 65, 101, 102], which might provide cognitive and/or physiological flexibility, and narrower-window entropies can relate to better performance [86] and faster orienting [103]. Fewer studies have examined the behavioral correlates of broader-window entropy, although several link this metric to greater awareness and consideration under harder conditions [86, 88, 93, 94]. Indeed, we predicted that higher broader-window entropy would relate to lower compulsive intake, with little relation to alcohol-only, based on our speculation that compulsion requires reduced attention to and awareness of negative consequences in order to keep drinking. In addition, β adrenergic receptors blockers decrease broader-but not narrower-window entropy in stressed rats, suggesting some functional dissociation [90]. Thus, considerable additional studies will be required to understand the physiological and behavioral importance of different forms of HRV entropy. Even so, our findings implicate higher broader-window entropy as negatively associated with compulsive consumption.

One important limitation is that the limited sample size of females strongly reduces our ability to make firm conclusions about whether there are actual sex differences, or just absence of evidence in one sex. Also, the selected HRV metrics may be better tuned to the male phenotype under this paradigm, and females often use different response strategies compared to males [46], which could influence the HRV-intake relationship. Indeed, females demonstrate greater persistence for alcohol [29] and more “addiction-like phenotypes” than males [104]. Our findings could also reflect sex differences in brain circuit activity related to HRV, as seen in humans [105–107]. Thus, future work is needed to clarify mechanisms underlying any HRV-drinking relations in females and males, including how sex hormones impact HRV-alcohol relations [66, 108].

Another important consideration is that we examined a single alcohol-drinking behavior, front-loading, since faster intake rates have been associated with great drinking problems in humans [54–57]. Even so, front-loading reflects just one of several other critical and clinically relevant aspects of alcohol behavior, including continued responding despite high intoxication, drinking promoted by alcohol cues, and the impacts of dependence, impulsivity, anxiety, and cognitive disfunction. Thus, it will be important to determine useful biomarkers for these other contributors in order to enhance alcohol treatment. For example, dependence models in rodents might better reflect the cardiovascular impacts of longer-term human drinking [109, 110], in parallel with any arousal patterns related to drive for alcohol. In addition, we focused on front-loading because it reflects an important and perhaps relatively stereotyped response pattern, which might illuminate HRV metrics associated with alcohol drives. However, front-loading is likely influenced by multiple, separable factors including anticipation, initiation, and motivation to consume, and our studies do not differentiate or shed light on these possibilities. In addition, our studies cannot determine whether intake-related HRV patterns were a consequence of the long drinking history, and future studies can examine HRV drinking metrics earlier in intake history. Thus, attempts to infer cognitive or behavioral strategies from HRV metrics remain highly speculative. We also note that HRV biomarker changes can occur but not correlate with behavior or subjective state [111, 112], making HRV metrics just one of several potential types of clinically-useful biomarkers.

In conclusion, by focusing on including non-linear and spectral HRV metrics, we found that combining such indicators can distinguish alcohol-only from compulsive intake with some sex differences. Thus, we provide novel HRV candidate biomarker patterns that could indicate individual and sex similarities and differences in drive for alcohol depending on the challenge level. Elucidating such possibilities will require additional future human studies. In addition, we hope our findings encourage others to examine HRV patterns in human drinkers and other drinking models to better understand the potential value of such metrics for alcohol treatment.

## DATA AVAILABILITY

Raw data is available from the corresponding author upon reasonable request.

## AUTHOR CONTRIBUTIONS

Dr. De Oliveira Sergio and Dr. Frasier made substantial contributions to the conception and design of the study, data collection, curation, and analyses, and writing and revising the manuscript. Mr. Zareb, Mrs. Wean, and Dr. Starski contributed substantially to data curation and analysis, interpretation of findings, and writing and revising the manuscript. Dr. Grippo contributed substantially to interpretation of findings and to critical revisions of the manuscript. Dr. Hopf made substantial contributions to the conception and design of the study, analysis and interpretation of findings, and drafting and revising the manuscript.

